# Towards more accurate preclinical glioblastoma modelling: reverse translation of clinical standard of care in a glioblastoma mouse model

**DOI:** 10.1101/2021.06.17.448792

**Authors:** Matteo Riva, Sien Bevers, Roxanne Wouters, Gitte Thirion, Katja Vandenbrande, Ann Vankerckhoven, Yani Berckmans, Jelle Verbeeck, Kim De Keersmaecker, An Coosemans

**Affiliations:** Laboratory of Tumor Immunology and Immunotherapy, Department of Oncology, Leuven Cancer Institute, KU Leuven, 3000 Leuven, Belgium; Department of Neurosurgery, Mont-Godinne Hospital, UCL Namur, 5530 Yvoir, Belgium; Oncoinvent, A.S., 0484 Oslo, Norway; Laboratory for Disease Mechanisms in Cancer, Department of Oncology, Leuven Cancer Institute, KU Leuven, 3000 Leuven, Belgium

**Keywords:** Glioblastoma, HGG, Mouse model, Preclinical, CT-2A, Surgery, Radiotherapy, Temozolomide, Combination therapies

## Abstract

**Introduction:** Contrarily to clinical trials, where glioblastoma (GBM) patients are usually heavily pre-treated (surgery, radiotherapy (RT) and temozolomide (TMZ)) before receiving the experimental therapies, in preclinical GBM studies experimental treatments have mostly been administered as monotherapy or combined with a very limited part of the clinical standard of care. This discrepancy can explain the failed clinical translation of preclinically successful treatments. This study is aimed at evaluating the feasibility and survival impact of the clinical standard of care in glioma-bearing mice. Methods. Neurospheres CT-2A tumor-bearing mice were treated with fluorescence-guided tumor resection, focal RT and oral TMZ.

**Results:** The implementation of postoperative intensive care treatment reduced surgical mortality by approximately 50% and increased experimental cost-effectiveness. Partial and total tumor resection significantly prolonged survival, the latter showing the strongest effect. Mice treated with surgery combined with RT or with RT and TMZ survived significantly longer than untreated controls.

**Conclusions:** Implementing the current GBM clinical standard of care in preclinical research is feasible and it leads to results in line with those obtained in patients. Systematic preclinical evaluation of the efficacy of experimental therapies in combination with standard of care treatments could improve the translational impact of next generation GBM animal studies.

## 1. Introduction

Glioblastoma (GBM) still represents an unmet medical need [1]. After diagnosis, GBM patients are treated with a combination of surgery, radiotherapy (RT) and chemotherapy (Temozolomide (TMZ)) [2].. However, the tumor almost invariably develops resistance mechanisms leading to the onset of recurrences with a median progression free survival (PFS) of seven months [3,4]. At this stage, none of the available therapeutic options is truly effective, leading to a median overall survival (OS) of GBM patients of 14 months [5,6].. Such dismal prognosis did not substantially changed in the last 15 years [1]. Despite the fact that many treatments have been proven to be effective in preclinical testing, none of them was capable to significantly prolong patients survival [7]. Even the most recent randomized clinical trials for checkpoints inhibitors showed negative results despite being supported by seemingly good preclinical data, confirming the presence of a significant translational gap between the preclinical setting and the clinical reality [8–10]. Such gap can be in part explained by the obvious differences between animal models and patients. Nevertheless, another important issue must be pointed out. Until now, in preclinical studies, the experimental therapies have been administered as a stand-alone therapy or, at best, in combination with only a very limited part of the multimodal clinical standard of care. In contrast, when preclinically successful therapies are translated into the clinic, patients have already been heavily pre-treated with surgery, RT and chemotherapy. Furthermore, in many cases, these patients also already developed tumor recurrence, which is nearly never the case in preclinical models, where treatment usually starts three to seven days after tumor inoculation. It has been shown that over time important modifications occur in gliomas and that recurrences show relevant differences when compared with their primary counterparts; furthermore, such evolutionary trajectory does not only depend on the characteristics of the original tumor, but also on the type of treatments that the patients receive [11,12]. In this context, it is clear that an untreated preclinical tumor model is not accurate enough to appropriately mimic the situation of heavily-pretreated human recurrent GBMs. Therefore, testing experimental treatments solely in such untreated animal models greatly undermines the translational value of their results. In order to overcome these limitations, we developed a preclinical research platform which allows to treat glioma-bearing mice with the complete human standard of care. Testing the efficacy of new experimental treatments in combination with fluorescence-guided surgery, focal RT and TMZ could increase the translational value of preclinical experiments and therefore the possibility of a successful clinical application.

## 2. Results

### The extent of tumor resection influences murine survival

In the preliminary experiment, NS/CT-2A tumor bearing mice were treated with surgery alone. Fluorescein-induced tumor fluorescence allowed to perform either a complete resection, in order to simulate clinical situations where the gross total removal of the tumor is possible, or an intentionally partial resection, in order to simulate clinical situations where the gross total removal of the tumor is not possible (due, for instance, to the invasion of eloquent brain areas). As shown in Figure 4, both partial and total tumor resection were capable to significantly prolong survival compared to controls (p<0.0001 and p=0.0011, respectively). Complete resection also induced a survival prolongation compared to partial resection, even if this difference did not reach statistical significance (p=0.0949). However, Logrank test for trend was statistically significant (χ2=13.78, p=0.0002).

**Figure 1.**
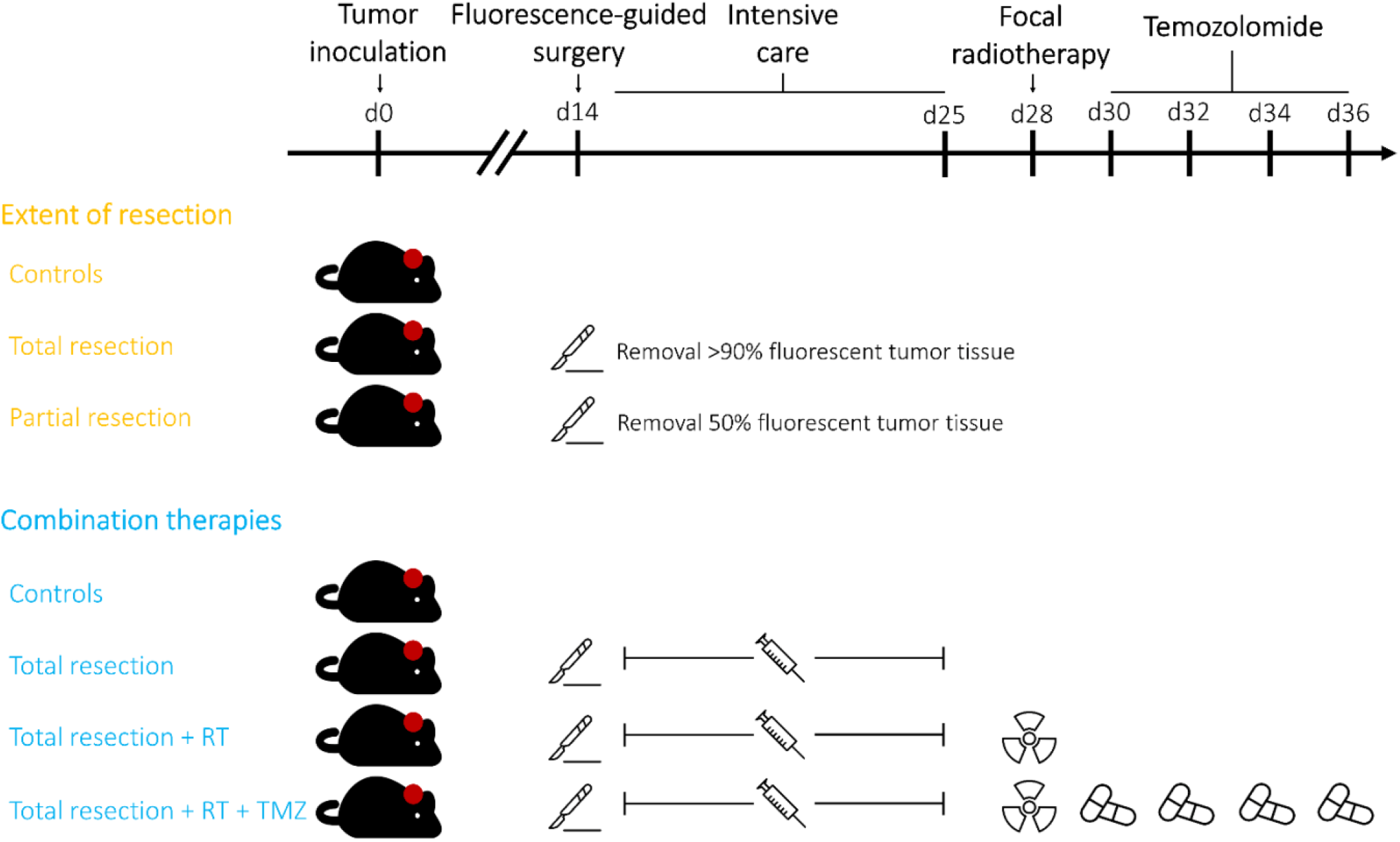
Experimental design. The preliminary experiment assessed the effect of the extent of surgical resection on murine survival. The combination experiment assessed the effect of tumor resection alone or in combination with RT and TMZ on survival. RT, radiotherapy; TMZ, Temozolomide.

**Figure 2.**
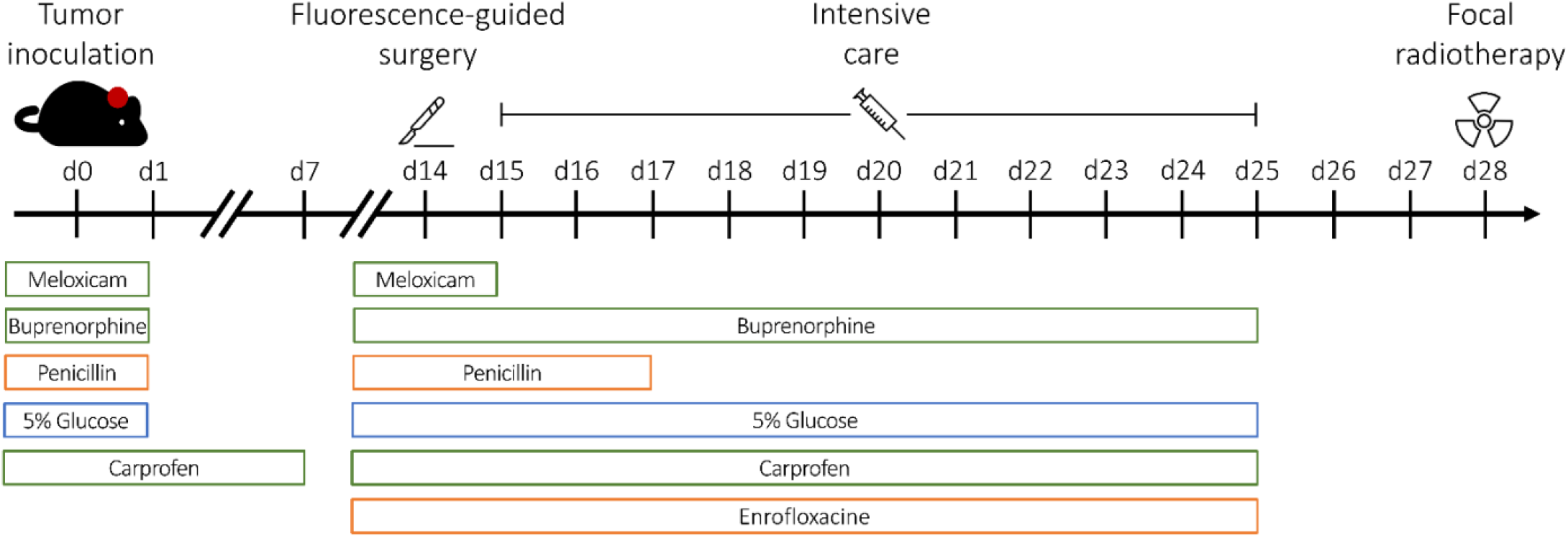
Timeline of drug administration. Timeline of the administration of antibiotics, pain medication and fluids following tumor cell inoculation and surgical removal of brain tumors.

**Figure 3.**
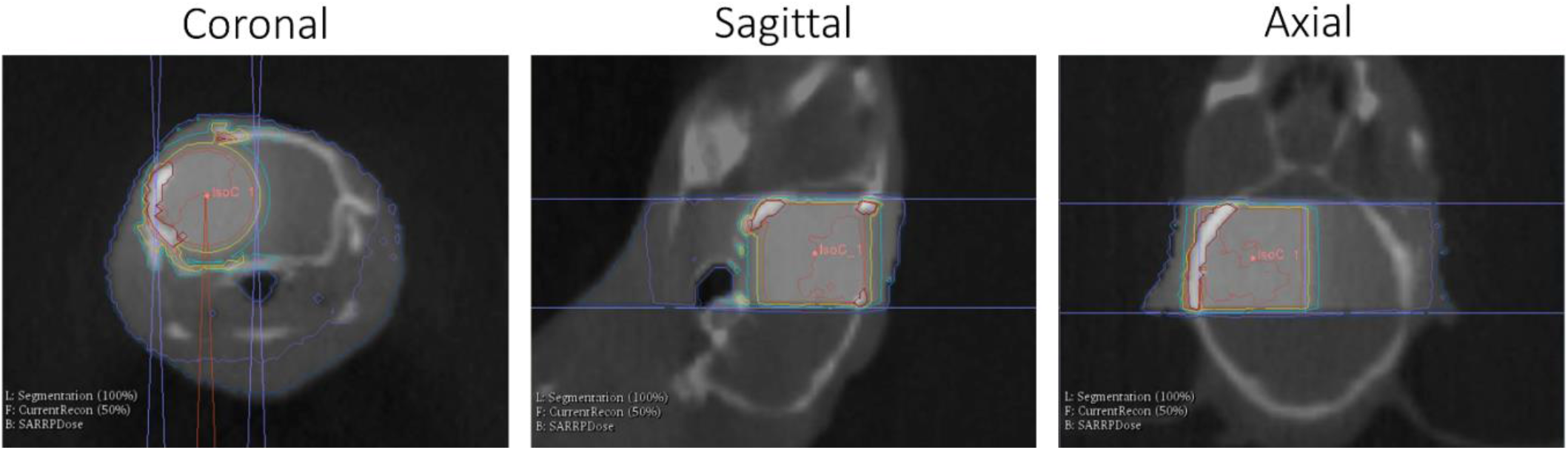
Treatment plan for focal radiotherapy. Isodose lines computed on the cone beam computed tomography (CBCT) indicate that the region of the surgical cavity and the surrounding infiltrated brain parenchyma receive the prescribed radiation dose.

**Figure 4.**
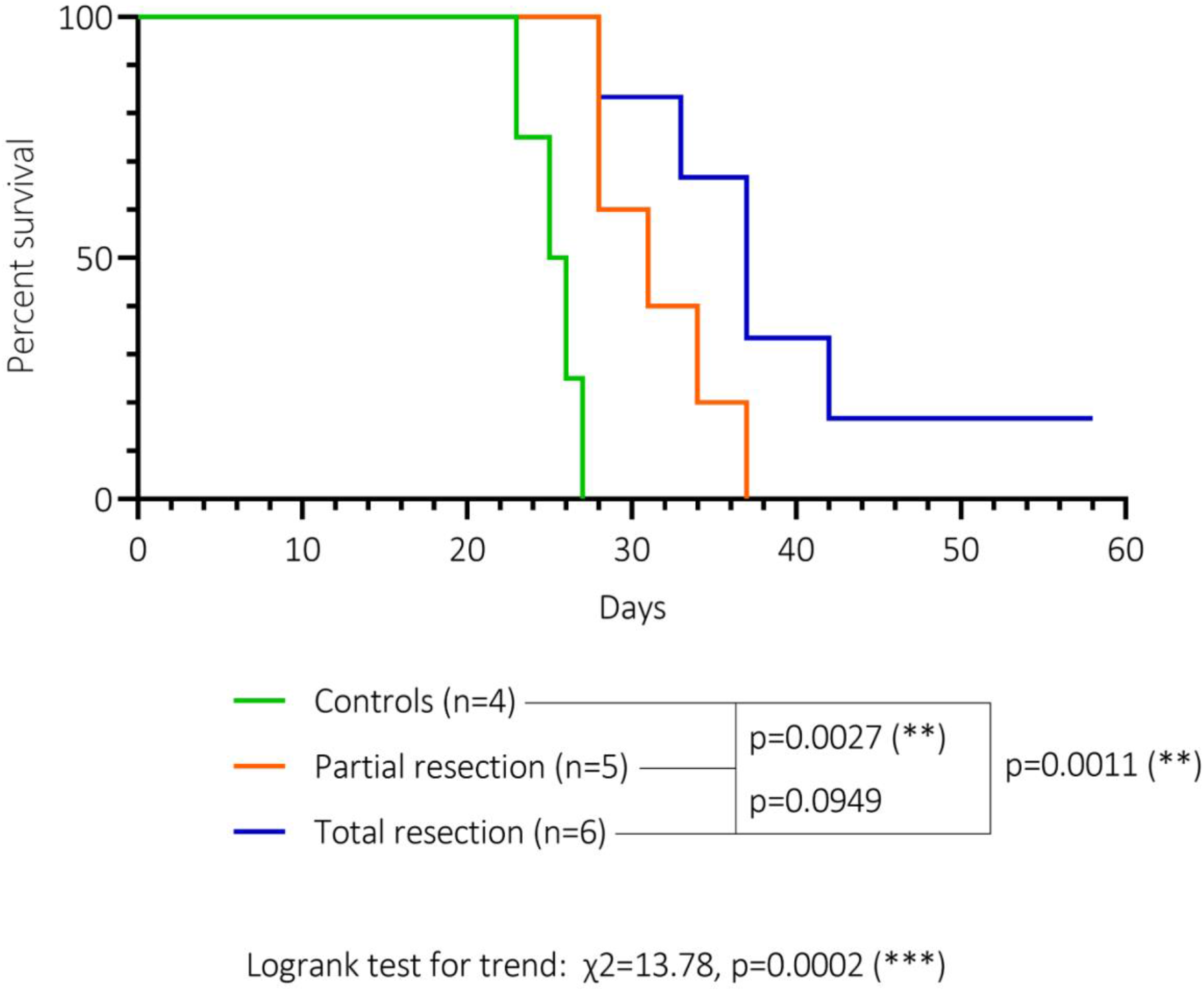
Effect of the extent of tumor resection on the survival of NS/CT-2A tumor bearing mice. Both mice undergoing partial and complete resection showed a significantly longer survival compared to controls (p=0.0027 and p=0.0011, respectively). Complete resection was able to prolong survival compared to partial resection; however, this difference did not reach statistical significance (p=0.0949). Logrank test for trend was statistically significant (χ2=13.78, p=0.0002).

### Effect of postoperative intensive care

Although rarely reported, surgical mortality represents one of the most important issues in performing tumor resection in glioma bearing mice. In our preliminary experiments, mice received only standard painkillers and antibiotics in the postoperative period. Under these conditions, approximately 60% of the mice died immediately after surgery or in the following days as a consequence of surgery itself. The implementation of the intensive care postoperative treatment protocol (as used in our combination experiment) lead to a reduction of perioperative mortality to approximately 30%.

### Sequential therapeutic steps induced a stepwise survival prolongation

Besides the above-mentioned surgical mortality, no adverse events or treatment related mortality were observed. The application of the full clinical standard of care to NS/CT-2A bearing mice was therefore proven to be feasible. As shown in Figure 5, mice treated with tumor resection combined with RT or with tumor resection combined with RT and TMZ survived significantly longer then controls (p=0.0423 and p=0.0022, respectively). Compared to controls (median survival 34 days), each sequential therapeutic step promoted a survival prolongation, even if these differences did not reach a statistical significance (median survival 47 days for surgical resection alone, not reached for surgery plus RT and surgery plus RT plus TMZ). However, Logrank test for trend was statistically significant (χ2=7.934, p=0.0049)

**Figure 5.**
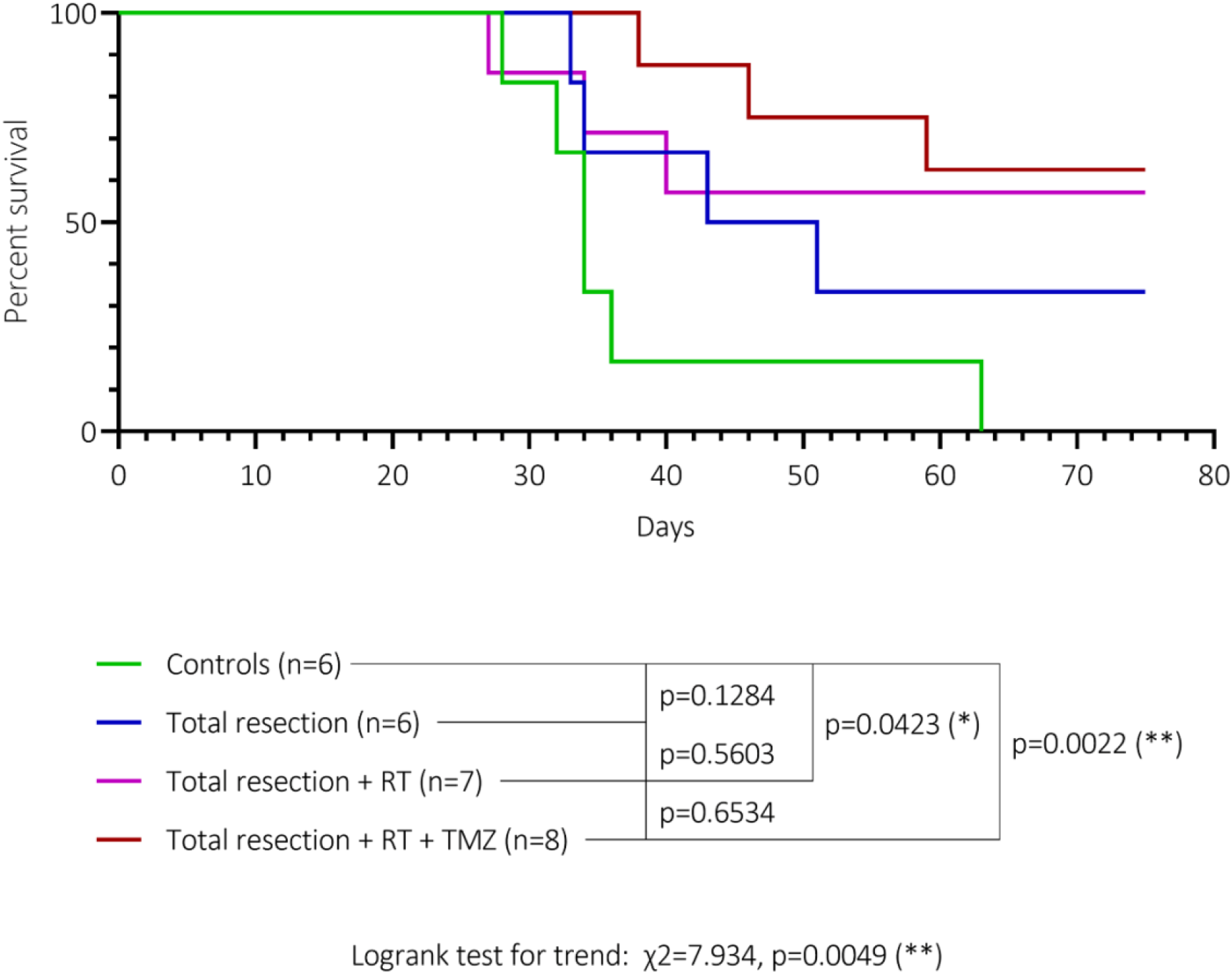
Effect of complete surgical removal, RT and TMZ on the survival of NS/CT-2A tumor bearing mice. Mice treated with surgery plus RT or surgery plus RT plus TMZ showed a significant survival prolongation compared to controls (p=0.0423 and p=0.0022). Each therapeutic step induced an improvement in survival. Even if direct comparison of controls vs surgery, surgery vs surgery + RT and surgery + RT vs surgery + RT + TMZ did not reach statistical significance (p=0.1284, p=0.5603 and p=0.6534; respectively), Logrank test for trend was statistically significant (χ2=7.934, p=0.0049).

## 3. Discussion

After the EORTC 26981–22981/NCIC CE3 trial, which demonstrated the efficacy of TMZ combined with RT in prolonging patients survival, all subsequent randomized studies assessing the efficacy of new treatments against high grade gliomas (HGG) invariably failed, despite being based on apparently good preclinical results [7]. The most recent case is that of the checkpoint inhibitor Nivolumab (anti programmed cell death protein 1 (PD1)): despite “successful” preclinical studies, three different randomized trials in patients were uncapable to demonstrate a survival benefit in the experimental arm compared to controls [8–10]. These failures underscore the important translational gap currently present between the preclinical research and the clinical setting [13]. As shown in a review published by our group, a strong research investment directed to the development of new and more accurate tumor models could be noted, in the attempt to generate murine HGGs that would be as close as possible to their human counterpart [14]. However, this represents only part of the problem and another important discrepancy must be pointed out. In preclinical studies, experimental treatments are administered very early during tumor development and almost always as stand-alone therapies. Conversely, when applied to patients, the same treatments are usually administered after surgical resection, RT and TMZ or even at tumor recurrence. It has been demonstrated that gliomas evolve over time, that treatments can strongly drive this evolution and that recurrent tumors are biologically different from their primary counterpart [11,12,15]. Therefore, it clearly appears that an untreated preclinical tumor model is not accurate enough to appropriately mimic the situation of recurrent human GBMs, that are heavily pre-treated with multimodal therapies. In this view, the integration of the full human standard of care in preclinical experiments represents a critical step in order to narrow the above mentioned translational gap.

In this study, we treated NS/CT-2A tumor bearing mice with a fluorescence-guided surgical resection of the tumor mass combined with focal RT and oral TMZ. Tumor resection alone or in combination with other treatments has been anecdotally implemented in the preclinical settings (some examples in [16–18]). However, to the best of our knowledge, this is the first study where fluorescence-guided surgical resection has been combined with focal RT targeted to the tumor cavity and its close surroundings and with oral TMZ. Treatments were started at day 14 after tumor cells inoculation, when a well-established tumor mass was present in accordance with the clinical situation at tumor diagnosis. It has been shown that the extent of resection (EOR) represents an independent prognostic factor for GBM, with patients undergoing gross total removal of the tumor surviving significantly longer than patients receiving partial resection [19,20]. In this study, we were able to model these two different situations and we confirmed their impact on survival. The possibility to study the response to treatments in these two scenarios is extremely important, since these two categories of patients could benefit from different therapeutic approaches. Although barely mentioned in the available literature, performing brain tumor resection in mice is challenging and the rate of fatal complications is high. In this study, we implemented for the first time an intensive care treatment protocol to support animals during the first two postoperative weeks. Following such protocol, we were able to reduce surgical mortality by 50%, thereby improving the ethical aspect and cost-effectiveness of our experiments. The analysis of the survival of mice undergoing surgical resection alone, combined with RT or combined with RT and TMZ shows that each supplementary therapeutic step promoted a further survival prolongation. This study therefore proves that implementing the clinical standard of care in preclinical GBM research is feasible and it induces results similar to those seen in patients. We believe that the systematic evaluation of the efficacy of experimental therapies in combination with the standard of care represents a fundamental prerequisite for the next generation of GBM preclinical studies. This will allow to confirm therapeutic efficacy in a scenario closer to clinical reality, to optimize the combination with standard of care treatment at the preclinical level and to reduce the risk of failure or clinical trials.

This study suffers from some limitations. First, the efficacy of surgical resection combined with RT or with RT and TMZ appears to be higher than what observed in patients [2]. This could partially limit the translational impact of our approach; therefore, we are currently optimizing our protocol in order to obtain results closer to the clinical reality. Second, we administered RT in one single dose and we did not perform concomitant chemotherapy, contrarily to the fractionated RT regimens associated with low-dose concomitant chemotherapy usually administered to patients [2]. As we already demonstrated by us and other groups, fractionated RT and concomitant TMZ in mice is feasible; therefore, we are currently implementing these therapies in our treatment pipeline [16,21]. Third, GBM patients are very often treated with corticosteroids in order to reduce brain edema. Analyzing the effect of corticosteroids in GBM is beyond the scope of this study; however, also in this case, we are currently implementing such treatment in our platform given the high clinical relevance [22]..

## 4. Materials and methods

### Experimental design

The experimental design is shown in Figure 1. Immunocompetent glioma-bearing mice were treated starting on day 14 after tumor cell inoculation, when a well-established tumor mass was present. In a preliminary experiment, we tested the efficacy of fluorescence-guided partial and complete tumor removal. In the combination experiment, mice were treated with fluorescence-guided complete tumor resection alone or followed by image-guided RT or by image-guided RT and oral TMZ. The triple combination of surgery, RT and TMZ represents the reverse translation in mice of the Stupp protocol currently in use as standard of care treatment for GBM patients [2].

### Tumor model

The orthotopic, immunocompetent NS/CT-2A glioma model recently developed by our group was used in this study [23]. CT-2A cells were provided by Prof. Thomas Seyfried (Boston College, Boston, MA) [24]. Following an 11-day protocol, CT-2A NS were generated in serum-free Dulbecco’s Modified Eagle Medium / Nutrient Mixture F-12 (DMEM/F-12, Sigma-Aldrich) supplemented with epidermal growth factor (EGF, Thermofisher), fibroblast growth factor (FGF, Thermofisher) and B27 supplement (Thermofisher). After enzymatical dissociation (Stempro Accutase, Thermofisher), 1×10^4^ NS-derived CT-2A cells were stereotactically inoculated in the right brain hemisphere of female, 12-14 week old C57BL/6 mice (Envigo), 2.5 mm lateral from the midline, 0.5 mm anterior to the bregma and 2.5 mm below the dura mater. During the whole procedure, the mice were kept under general anesthesia with an IP injection of 6μl/g body weight of a mixture of 18.75mg/ml ketamine (Pfizer) and 0.125% xylazine hydrochloride (Bayer). During the perioperative period, mice received antibiotics (penicillin IP, Kela Pharma), painkillers (meloxicam (Boheringer) and buprenorphine (Ceva) IP, carprofen (Zoetis) in drinking water) and fluids (5% glucose solution IP, Baxter) to compensate possible blood loss (Table 1, Figure 2).

**Table 1.** Perioperative drugs for tumor inoculation and tumor resection.

|  | Drug | Dosage | Administration route | Duration |
| --- | --- | --- | --- | --- |
| Tumor inoculation | Penicillin | 100.000 UI/Kg | IP | 1x before tumor inoculation |
|  | 5% Glucose | 1ml/mouse | IP | 1x before tumor inoculation |
|  | Meloxicam | 5 mg/kg | IP | 1x after tumor inoculation |
|  | Buprenorphine | 0.1 mg/kg | IP | 1x after tumor inoculation |
|  | Carprofen | 0.067 mg/mL | Drinking water | 7 postoperative days |
| Tumor resection | Penicillin | 100.000 UI/Kg | IP | 1x before tumor resection, twice daily for 3 postoperative days |
|  | 5% Glucose | 1ml/mouse | IP | 1x before tumor resection, twice daily for 7-10 postoperative days (until complete recovery) |
|  | Meloxicam | 5 mg/kg | IP | 1x after tumor resection |
|  | Buprenorphine | 0.1 mg/kg | IP | 1x after tumor resection, twice daily for 7-10 postoperative days (until complete recovery) |
|  | Carprofen | 0.067 mg/mL | Drinking water | 14 postoperative days |
|  | Enrofloxacin | 0.25mg/ml | Drinking water | 14 postoperative days |

IP: intraperitoneally

### Fluoresce-guided microsurgical resection of brain tumors

The surgical removal of brain tumors was performed on day 14 after tumor cell inoculation. To induce fluorescence in the brain tumors, mice received with 5mg/kg fluorescein (Sigma) dissolved in 100µl of 0.9% NaCl solution (Braun) via tail vein injection immediately before surgery, as previously described [16]. Mice were then placed under general anesthesia with 2% isoflurane (Baxter), administered with penicillin and 5% glucose solution IP (Table 1, Figure 2) and transferred on a stereotactic frame (Stoelting) below the surgical microscope (Leica Microsystems). This microscope can be used in white light or in ultra violet light combined with a green fluorescent protein (GFP) filter, therefore making it possible to carry out surgical procedures both in standard conditions and by visualizing the fluorescein-induced tumor fluorescence. After disinfecting the skin with Isobetadine (Mylan), the midline head incision used to inoculate tumor cells was reopened. Under microscopic view and in white light, a round craniotomy of approximately 4 mm in diameter and centered on the burr hole used for tumor cell inoculation was obtained using high-speed drill (Stoelting) equipped with a 2 mm diamond ball-head. The dura mater was gently opened with fine forceps and irrigation with sterile 0.9% NaCl solution was used to control superficial bleedings. The microscope was then switched to UV light and the fluorescent tumor tissue was removed by means of a cell culture suction pump (Curis) connected to sterile 200µl tips. Bleeding from the brain tissue was controlled by irrigation with sterile 0.9% NaCl solution and by applying the hemostatic agent Floseal (Baxter). After disinfection, the skin was closed with 4-0 silk stitches. Mice were removed from the stereotactic frame, injected with meloxicam and buprenorphine IP and placed in warmed caged for postoperative recovery (Table 1, Figure 2).

### Postoperative intensive care treatment

An intensive treatment (Table 1, Figure 2) was carried 11 days postoperatively in order to facilitate mice recovery, reduce surgical mortality and therefore maximize the experimental output. During this 11-days period, mice had full access to soft high-caloric food (DietGel Boost, Clear H2O) and for the first five days the cages were kept warm by means of single-use heating pads (The Heat Company). Penicillin IP twice daily and enrofloxacin (Bayer) in drinking water were administered for three and 11 days postoperatively, respectively. Buprenorphine twice daily and 5% glucose solution once daily were administered IP for seven to 11 days postoperatively, until complete recovery. When operated mice reached the endpoints (see below) during the first two postoperative weeks, this was considered as surgical- and not as tumor-related and these mice were excluded from survival analysis.

### Radiotherapy

Mice that successfully recovered from surgery received focal radiotherapy 14 days after surgery as already described by our group [21,25]. A stereotactic, cone beam computed tomography (CBCT)-guided arc irradiation was performed using a Small Animal Radiation Research Platform (SAARP, Xstrahl Life Sciences). Mice were put under general anesthesia with isoflurane at 2% and transferred to the SAARP workstation, where the brain CBCT was acquired. The irradiation isocenter was centered on the craniotomy and placed 2 mm below the brain surface, in a position corresponding to the center of the surgical cavity. The treatment plan consisted of one 356° arc with a choice of a square collimator of 5×5 mm such that the entire surgical cavity received the prescribed dose of 4Gy in one single dose. Isodose lines confirmed that the region of the surgical cavity received the full radiation dose whereas the surrounding healthy brain was spared (Figure 3).

### Chemotherapy

Temozolomide capsules of 20mg (Temodal®, Merk Sharp & Dohme) were dissolved in 0.9% NaCl at a concentration of 10mg/ml. 10mg/ml of L-histidine (Sigma Aldrich) was added to the suspension to increase solubility. After 30 minutes in a 37 °C sonication bath, the temozolomide suspension was administered by oral gavage. Each TMZ-treated mouse received four doses of 50mg/kg on alternate days [25].

### Endpoints

Humane endpoints were used in this study. After the inoculation of tumor cells, mice were followed up for weight loss and development of clinical symptoms [26]. General endpoints for mice euthanasia during the study were the presence of grade III symptoms (hunched posture, severe hemi-paresis (constant circling behavior, frequent drops, inability to move) or both) or a weight loss higher than 20% compared to the day of tumor cell inoculation. However, pilot experiments (data not shown) demonstrated that in the postoperative period, our intensive care treatment allows mice to sustain and recover from weight losses greater than 20%. For this reason, during 11 days postoperatively a maximal weight loss of 40% was allowed if mice were not showing neurological symptoms. Mice that at the end of this period showed a weight loss ≥ 20% compared to the day of tumor cell inoculation were euthanized. Mice that at the end of the two postoperative weeks showed no weight loss or a weight loss ≤ 20% compared to the day of tumor cell inoculation remained in experiment and were randomized for further treatments.

### Statistics

Statistical analysis was performed with Prism v.9.0 (Graphpad Software). For survival data, a Logrank test was used and p<0.05 was considered significant.

### Ethics

All animal experiments were approved by the Animal Ethics Committee of the Katholieke Universiteit Leuven (project 109/2020 and 086/2019) which follows the Directive 2010/63/EU and the Belgian Royal Decree of 29 May 2013.

## 5. Conclusions

Implementing the full GBM clinical standard of care in preclinical research is feasible and induces results in line with the clinical reality. We believe that systematically evaluating the efficacy of experimental therapies in combination with tumor resection, RT and TMZ could improve the translational impact of next generation GBM animal studies and reduce the current gap between the preclinical setting and the clinical reality.

## Authors’ contribution

Conception of the study: MR, AC; Design of the experiments: MR, AC; In vivo experiments: MR, SB, RW, GT, KV, JV; Mouse follow-up: MR, SB, RW, GT, KV, AV, YB; Analysis and interpretation of the results: MR, AC; Manuscript writing: MR; Manuscript proof reading: SB, RW, GT, KV, AV, YB, JV, KDM, AC

## Data availability

Data supporting this study are available from the corresponding author upon request.

